# A View of Waste Might Decrease Relaxation: The Effects of Viewing an Open Dump in a Forest Environment on the Psychological Response of Healthy Young Adults

**DOI:** 10.1101/2020.08.19.256990

**Authors:** Bielinis Ernest, Janeczko Emilia, Takayama Norimasa, Słupska Alicja, Korcz Natalia, Zawadzka Anna, Bielinis Lidia

**Affiliations:** Department of Forestry and Forest Ecology, Faculty of Environmental Management and Agriculture, University of Warmia and Mazury, Pl. Łódzki 2, 10-727 Olsztyn, Poland; Department of Forest Utilization, Institute of Forest Sciences, University of Life Sciences in Warsaw, Nowoursynowska 159, Warsaw, Poland; Environmental Planning Laboratory, Department of Forest Management, Forestry and Forest Products Research Institute in Japan, 1 Matsunosato, Tsukuba, Ibaraki, Japan; Department of Natural Foundations of Forestry, Institute of Soil Science and Environment Management, University of Life Sciences in Lublin, Akademicka 13, Lublin, Poland; Department of Social Pedagogy and Methodology of Educational Research, Faculty of Social Science, ul. Żołnierska 14, 10-561 Olsztyn, Poland

## Abstract

Forest recreation can be successfully used for the psychological relaxation of respondents and can be used as a remedy for common problems with stress. The special form of forest recreation intended for restoration is forest bathing. These activities might be distracted by some factors, such as viewing buildings in the forest or using a computer in nature, which interrupt psychological relaxation. One factor that might interrupt psychological relaxation is the occurrence of an open dump in the forest during an outdoor experience. To test the hypothesis that an open dump might decrease psychological relaxation, a case study was planned that used a randomized, controlled crossover design. For this purpose, two groups of healthy young adults viewed a control forest or a forest with an open dump in reverse order and filled in psychological questionnaires after each stimulus. A pretest was used. Participants wore oblique eye patches to stop their visual stimulation before the experimental stimulation, and the physical environment was monitored. The results were analyzed using the two-way repeated measures ANOVA. The measured negative psychological indicators significantly increased after viewing the forest with waste, and the five indicators of the Profile of Mood States increased: Tension-Anxiety, Depression-Dejection, Anger-Hostility, Fatigue, and Confusion. In addition, the negative aspect of the Positive and Negative Affect Schedule increased in comparison to the control and pretest. The measured positive indicators significantly decreased after viewing the forest with waste, the positive aspect of the Positive and Negative Affect Schedule decreased, and the Restorative Outcome Scale and Subjective Vitality scores decreased (in comparison to the control and pretest). The occurrence of an open dump in the forest might interrupt a normal restorative experience in the forest by reducing psychological relaxation. Nevertheless, the mechanism of these relevancies is not known, and thus, it will be further investigated. In addition, in a future study, the size of the impact of these open dumps on normal everyday experiences should be investigated. It is proposed that different mechanisms might be responsible for these reactions; however, the aim of this manuscript is to only measure this reaction. The identified psychological reasons for these mechanisms can be assessed in further studies.

## Introduction

Forest recreation is an activity engaged in for pleasure, and it is done outside in a natural forest environment [1,2]. Recreation in a forest environment is a common activity of many people living in forested areas around the world. A good example is Poland, where the half of the surveyed adults declare that they participated in some form of forest recreation [3]. Forest recreation has an evidence-based, positive effect on the psychological and physiological relaxation of participants, who may suffer from anxiety, stress and other problems caused by living in a modern, urbanized environment [4]. This outdoor activity in a forest environment is often conceptualized as ‘forest bathing’ (if the aim of the activity is restoration) [5,6] or ‘forest therapy’ (if the aim of the activity is usage of the forest for healing) [7]. These activities unquestionably induce psychological relaxation [8–15] and have a positive influence on mental health [7,16–20]. Induced psychological relaxation might be measured using psychometric techniques. The most popular measured indices are the following: mood states, positive and negative affect, restorativeness and vitality [1,14,21–24].

During forest recreation, an undisturbed forest environment is important to produce the most valuable, most restorative experience, which is valued by the respondents involved in this recreation [25–27]. Some research suggests that some ‘disturbing factors’ that occur during nature recreation, like using a computer during this activity or seeing an urban building in the background of the forest landscape, might damage the recreational experience of respondents. This might be interpreted as a reduction of the restoration and psychological relaxation in comparison to an undisturbed control [28,29].

A common problem in some parts of the world, and also in Poland, is the illegal dumping of waste in a forest environment [30–32]. A dump occurring in a forest is a kind of ‘open dump’ [33] and they usually occur near urbanized areas where forest recreation is highly important [34]. That situation might influence the quality of the natural forest experience during recreation in urban and semi-urban forests because respondents might view dumps in the forest and that can influence their level of psychological relaxation. Thus, wastes are ‘disturbing factors’ in the forest environment. Besides, there are some examples of ‘disturbing factors’ that occur during recreational experiences described in literature; however, there is nothing known about the effect of an illegal dump in a forest environment on the mood, affect, restorativeness and vitality of respondents experiencing and viewing this disturbance.

Therefore, the aim of this study is to clarify whether the view of an open dump in a forest environment (in comparison to an undisturbed, control forest) might influence the psychological relaxation of respondents, and to clarify whether this view has a negative or neutral effect on respondents in a randomized, controlled crossover experiment.

## Materials and Methods

### Ethical statement

This study was ethically reviewed and approved by the Ethical Review Board of the University of Warmia and Mazury in Olsztyn. The number of this ethic statement is the following: 07/2019. All procedures performed in this study were conducted in accordance with the ethical standards of the Polish Committee of Ethics in Science and with the 1964 Helsinki Declaration and its later amendments.

### Participants

Twenty-four healthy participants were involved in this study (mean age 21.63 years ± 1.18 SD, 18 males and 6 females). The participants were young adults studying at the University of Warmia and Mazury in Olsztyn who had graduated high school and had no regular working activity. Only healthy participants of Polish nationality without any history of mental or physical illness were involved. None were taking any medication that could influence the psychological or physiological indices. The participants received a short description of the research before the experiment with information on the details of the experiment and without informing them about the expected results. All participants’ consent to participate in the experiment and their involvement was voluntary.

The participants were randomly divided into two groups, group A and group B (twelve persons in each). Group A first participated in the stimulation of an undisturbed forest and after 15 minutes of rest participated in the stimulation with the disturbed forest. Group B participated in this activity in the reverse order according to the crossover design (Table 1).

**Table 1.** Procedure followed during the experiment

| Date | Group A (n = 12) |  | Group B (n = 12) |  |
| --- | --- | --- | --- | --- |
| 2019/06/14 | Time | Activity | Time | Activity |
|  | 10:00 | Meeting at the gathering point | 13:00 | Meeting at the gathering point |
|  | 10:15 | 15 minute walk into the forest stand | 13:15 | 15 minute walk into the forest stand |
|  | 10:30 | 15 minutes with eye-patches | 13:30 | 15-minutes with eye-patches |
|  | 10:45 | Filling in the questionnaires (pretest) | 13:45 | Filling in the questionnaires (pretest) |
|  | 11:00 | Viewing the control forest | 14:00 | Viewing the forest with open waste |
|  | 11:15 | Filling in the questionnaires (posttest) | 14:15 | Filling in the questionnaires (posttest) |
|  | 11:30 | 15 minutes with eye-patches | 14:30 | 15 minutes with eye-patches |
|  | 11:45 | Filling in the questionnaires (pretest) | 14:45 | Filling in the questionnaires (pretest) |
|  | 12:00 | Viewing the forest with open waste | 15:00 | Viewing control forest |
|  | 12:15 | Filling in the questionnaires (posttest) | 15:15 | Filling in the questionnaires (posttest) |
|  | 12:30 | End of the experiment | 15:30 | End of the experiment |

### Experimental stimuli

During the experiment, each participant was involved in two kinds of stimulating activities (posttests) or non-stimulating activities (pretests) in a standing position. During these activities, talking and using electronic devices was not allowed. Also, the participants stood in a line and did not touch each other. While standing, small movements were allowed. Natural standing was appreciated and also sitting on the litter was allowed to simulate a more natural forest recreational experience. The participants were required to look exactly at the forest litter from one meter away or look at the open dump during the stimulation. While completing the questionnaires, participants were required to look only at their own paper questionnaire and looking around was not allowed. To measure the states of participants without stimulation in the experiment (stimulation was induced by a disturbed or undisturbed forest environment), black opaque eye patches were administered. The participants spend 15 minutes in each forest setting using the eye patches before the measurement. During, this time the participants did not view each forest environment, and so visual stimulation was not induced. The participants standing in the pretest were a few meters from the open dump to avoid being stimulated by the smell. After that time, the participants took off the patches, stepped closer to the environmental settings and viewed the undisturbed or disturbed forest environment for another 15 minutes to induce visual stimulation. The undisturbed (control) forest environment was a broad-leaved suburban forest in the vegetation stage located near the city of Olsztyn in northeast Poland. The composition of the matured tress (medium age: 90 years) was the following: white oak 80%, white lime 10%, and black alder 10% (Figure 1A). The disturbed forest environment (experimental variant) was located 50 meters from the control forest with a similar composition of tree species and ages. The disturbance in the forest was an open dump with plastic bottles, glass bottles, cans and others small waste (Figure 1B) on the ground. In both forest environments, there was the presence of hornbeam seedlings in the undergrowth and old leaves on the litter. The season of the experiment was summer.

**Figure 1.**
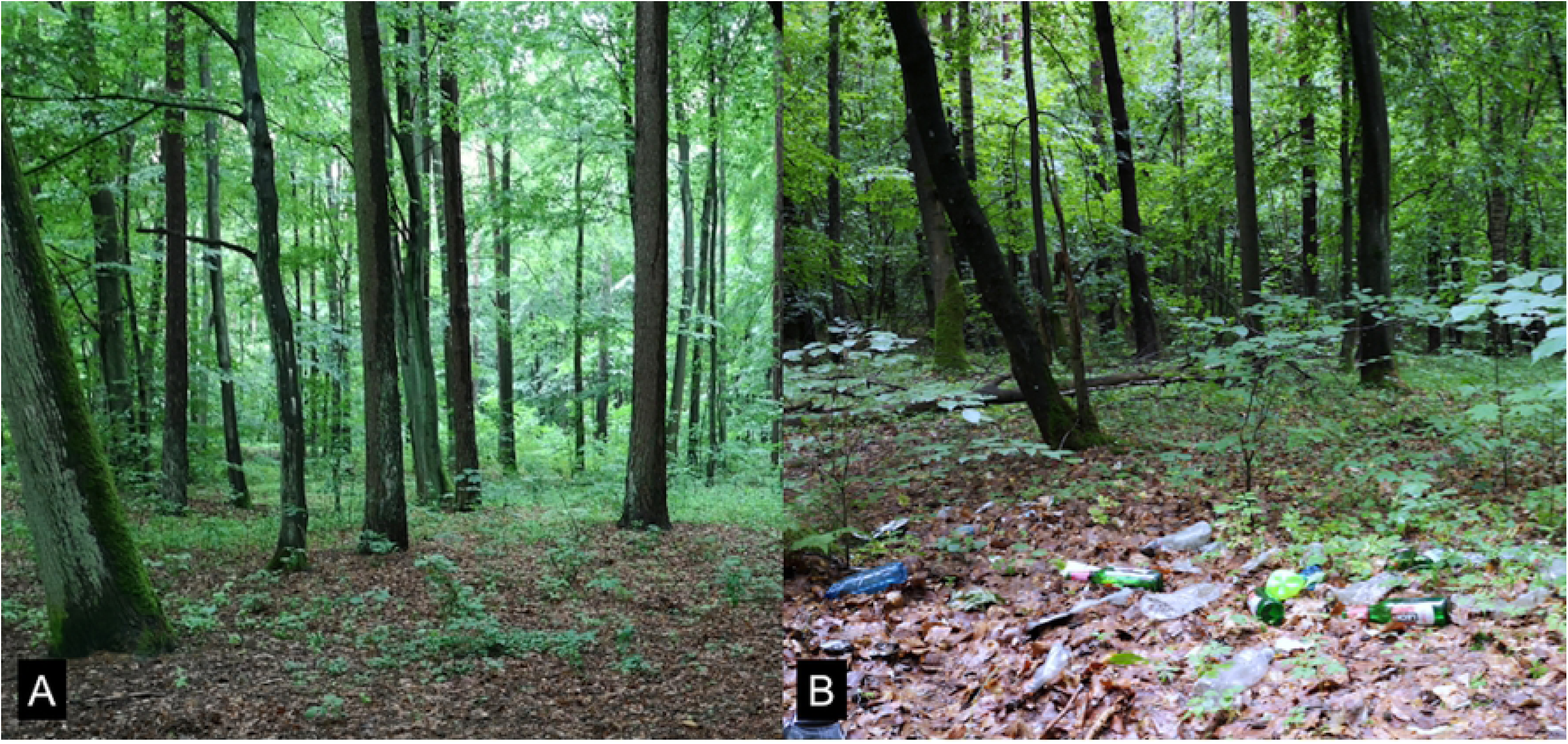
Photos showing the control forest (A) and the forest with open dump (B).

### Procedure

Each participant was involved in the measurements four times: i) before viewing the undisturbed forest environment, ii) after viewing the undisturbed forest environment, iii) before viewing the disturbed forest environment, and iv) after viewing the forest environment. The order depended on whether they belonged to group A or group B (Table 1). The psychometric questionnaires were administered for participants after each time, and each time the participants filled in the following: Profile of Mood States (POMS), Positive and Negative Affect Schedule (PANAS), Restorative Outcome Scale (ROS), and Subjective Vitality Scale (SVS).

### Measurements

In this research, four different psychological questionnaires were used. The Profile of Mood States is a commonly used questionnaire for measuring six different mood states: tension-anxiety, depression-dejection, anger-hostility, fatigue, confusion and vigor. The questionnaire is valid and commonly used [35]. In this research, the Polish version with 65 items using a 4-point Likert scale was applied. The Positive and Negative Affect Schedule (PANAS) is also a commonly used questionnaire that measures two kinds of emotional affect: positive and negative. This questionnaire, containing 20 items, is valid and reliable [36], and also the Polish version using a 5-point Likert scale was applied. The Restorative Outcome Scale measured the restorative effect of each environment. It contained six items, and the scale has been found to be valid and reliable [37]. The Subjective Vitality Scale contained four items that measured vitality, and the scale has been found to be valid and reliable [38]. For both these scales, the Polish version using a 7-point Likert scale was applied.

A previous study showed that all these questionnaires have moderate (Cronbach’s α = 0.794) to high (Cronbach’s α = 0.921) reliability in case of young Polish adults and might be used with adequate precision in this study [1].

During the experiment, the values describing the physical environment were measured. In the control forest environment and in the forest with the open dump, these values were recorded ten times for each environment using commercially available meteorological tools (temperature and humidity) and smartphone applications (illuminance and sound pressure).

### Data and statistical analysis

The raw data from the questionnaires were used for statistical analysis. The data were expressed as the mean values ± standard deviation. Two-way repeated measures ANOVAs were conducted for the analysis; and the effects of ‘time’, ‘conditon’ and the interaction ‘time × condition’ were analysed using the results for the POMS, PANAS, ROS and SVS scales. SigmaPlot 12.0 for Windows (Systat Software, Erkrath, Germany) was used for the ANOVA calculations. The SPSS 25 Software for Mac (IBM, Armonk, NY, USA) was used for the t-test calculations.

The statistical power analysis was conducted using the free G*Power 3.1.9.4 software for Mac (Heinrich Hein University, Düsseldorf, Germany) [39]. The actual power (1–β error probability) was calculated as 0.816. The ‘ANOVA: repeated measure, within factors’ statistical test was used, and a power analysis type of ‘Post hoc: Compute achieved power’ was applied with a sample size of 0.25 and an α probability error of 0.05. Usually, the power of proper prepared experiments is 0.8 or higher, and so the statistical power in this experiment is acceptable.

## Results

### Physical environment

Table 2 shows the results of the physical environment measurements for the control (forest without waste) and experimental (forest with waste) conditions. Only the relative humidity was significantly higher in the forest with waste than in the control forest. Other physical measurements (temperature, illuminance, and sound pressure) did not differ significantly considering these two conditions.

**Table 2.** Results of the t-test comparison between the control and experimental settings (physical environment).

| Item | Control |  | Waste |  | *t* | *p* | *R* |
| --- | --- | --- | --- | --- | --- | --- | --- |
|  | Mean | SD | Mean | SD |  |  |  |
| Temperature (°C) | 20,71 | 1,36 | 21,22 | 1,11 | -0,737 | 0,47 | 0,907 |
| Relative humidity (%) | 73,6 | 2,22 | 77,5 | 3,14 | -3,209 | 0,005 | ** 0,957 |
| Illuminance (lux) | 3367,30 | 296,39 | 3073,80 | 623,74 | 1,344 | 0,196 | 0,04 |
| Sound pressure (dB) | 35,84 | 3,09 | 37,05 | 1,682 | -1,085 | 0,292 | 0,671 |

n = 10, \*\* p < 0.01

### Profile of Mood States

A two-way repeated measures ANOVA of the POMS values was conducted, with condition and time as the two factors (Table 3). The results showed that interactions occurred in the case of the five subscales of the POMS: Tension-Anxiety, Depression-Dejection, Anger-Hostility, Fatigue, and Confusion. In the case of the Vigor subscale, there was no statistically significant interaction. Main effects were statistically significant for the five subscales of POMS except Vigor. The results of multiple comparisons tests showed that (Table 4) there were no differences between the control before and after viewing the forest stand (Pre vs. Post) for all six subscales of the POMS. There were also no differences between the measures before viewing the control forest and the forest with waste (Pre: Control vs. Waste) for all six subscales of POMS. Furthermore, a comparison of the results of the post-hoc tests for ‘Waste: Pre vs. Post’ showed that after viewing the forest with waste, the participants had significantly higher values of the five negative mood indicators with the exception being Vigor, indicator of a positive mood, which was non-significant. There were also differences after viewing the control forest and the forest with waste after the experimental stimuli (Post: Control vs. Waste). The five negative mood indicators had significantly higher values after the stimuli with the forest with waste than that with the forest except for the positive indicator ‘Vigor’ (non-significant).

**Table 3.**
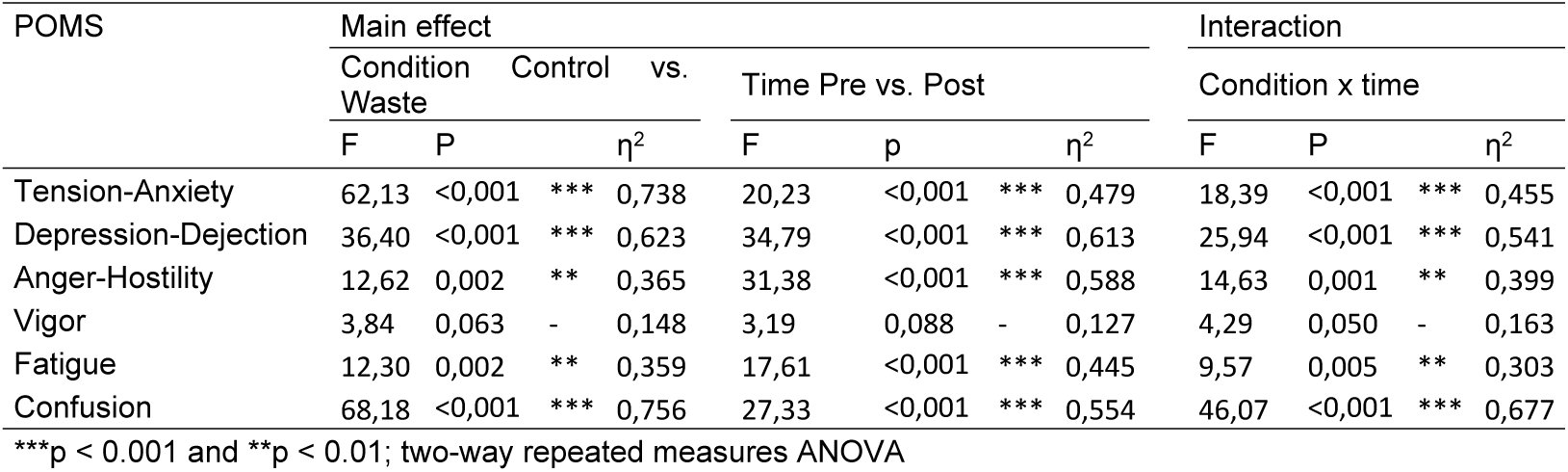
Results of two-way repeated measures ANOVA for the Profile of Mood States (mood).

**Table 4.**
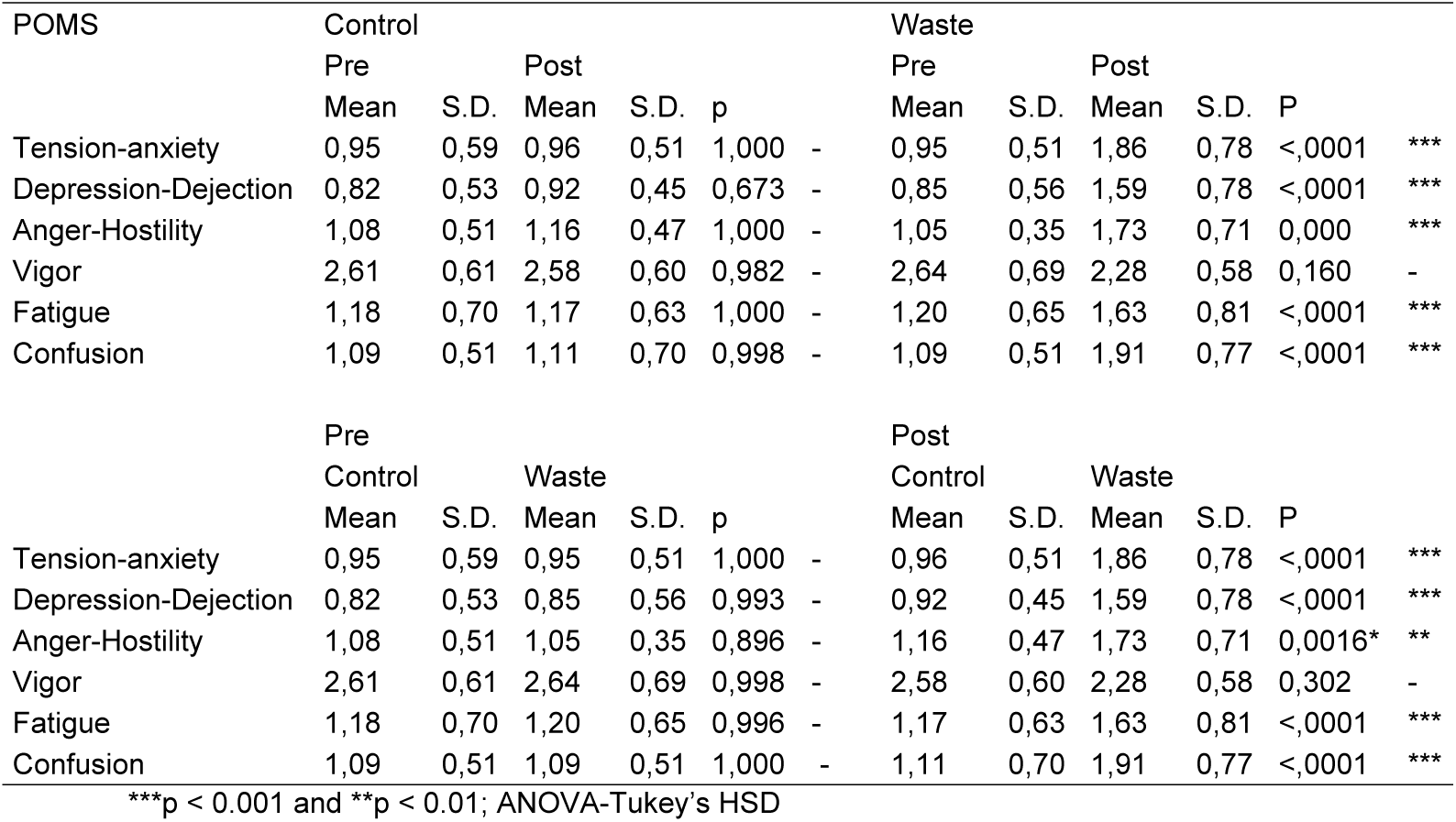
Results of the multiple comparison test between the control and experimental (setting) and pre-post (exposure to viewing) for the Profile of Mood States (mood).

### Positive and Negative Affect Schedule

The two-way repeated measures ANOVA was conducted to compare the differences in the PANAS Negative and PANAS Positive scores and to analyze the interaction between the factors and main effects (Table 5). Significant interactions were observed for both the ‘PANAS Positive’ and ‘PANAS Negative’ scores, and also all main effects were significant. The results of Tukey’s HSD comparisons showed (Table 6) that both PANAS indicators did not differ significantly before and after viewing the control forest environment (Control: Pre vs. Post) and before the experimental stimuli of the control forest or the forest with waste (Pre: Control vs. Waste). Also, this was true after viewing the forest environment with waste (Waste: Pre vs. Post). The positive aspect of PANAS decreased significantly, and the negative aspects of PANAS increased significantly. Furthermore, there were significant differences between the stimulation by the control forest and the forest with waste (Post: Control vs. Waste). The scores of the Positive aspect of PANAS were significantly lower after viewing the forest with waste, and the scores of the negative aspects of PANAS were significantly higher after the viewing forest with waste (both in comparison to the control).

**Table 5.**
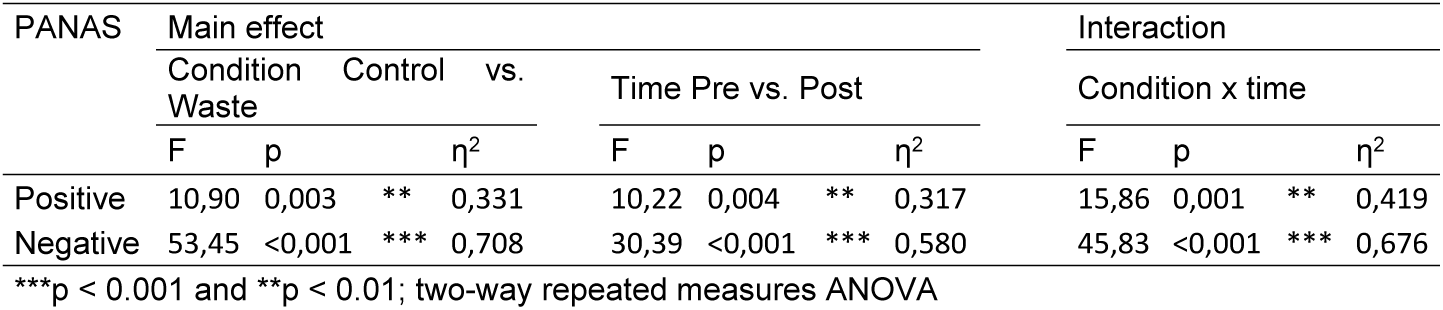
Results of two-way repeated measures ANOVA for Positive and Negative Affect Schedule (emotion).

**Table 6.**
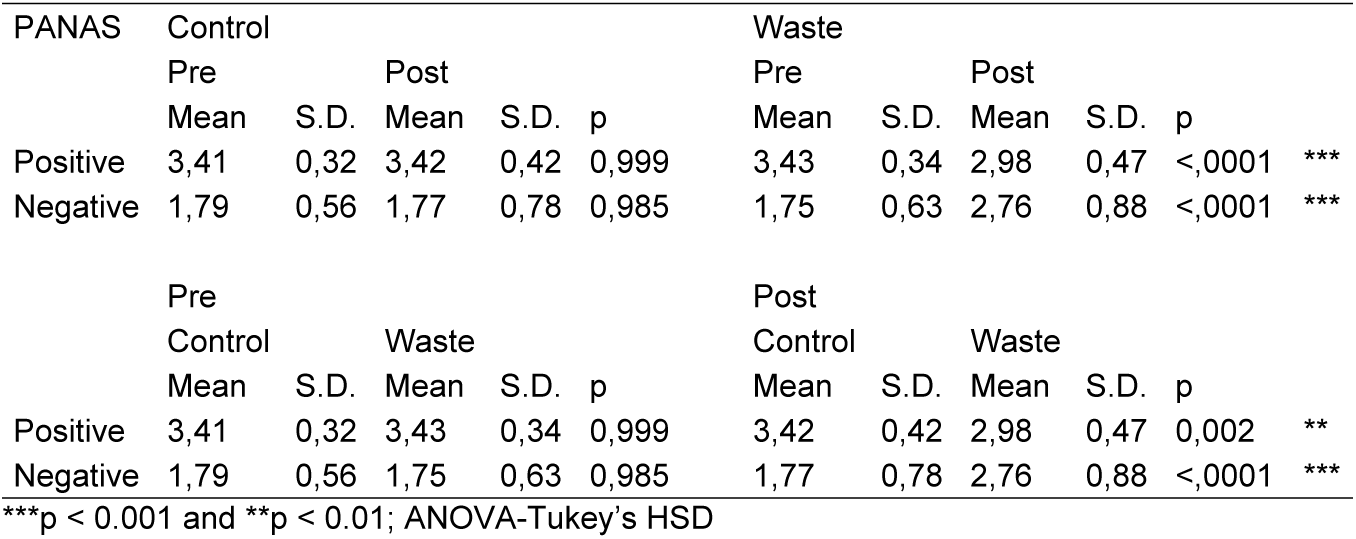
Results of a multiple comparison test between the control and experimental (setting) and pre-post (exposure to viewing forest) for the Positive and Negative Affect Schedule (emotion).

### Restorative Outcome Scale

Two factors ‘Conditon’ and ‘Time’ were analyzed using a two-way repeated measures ANOVA to compare the changes in the ROS scores and to analyze the interaction between factors (Table 7). Both main effects and also the interaction were significant. The results of Tukey’s HSD comparisons (Table 8) showed that ROS scores were not significantly different in both times in the control environment (Control: Pre vs. Post) and not significantly different before and after viewing the forest with waste (Waste: Pre vs. Post). In contrast, the ROS significantly decreased after viewing the forest with waste in comparison to the pre-test (Waste: Pre vs. Post) and significantly decreased after viewing the forest with waste in comparison to the control (Post: Control vs. Waste).

**Table 7.**
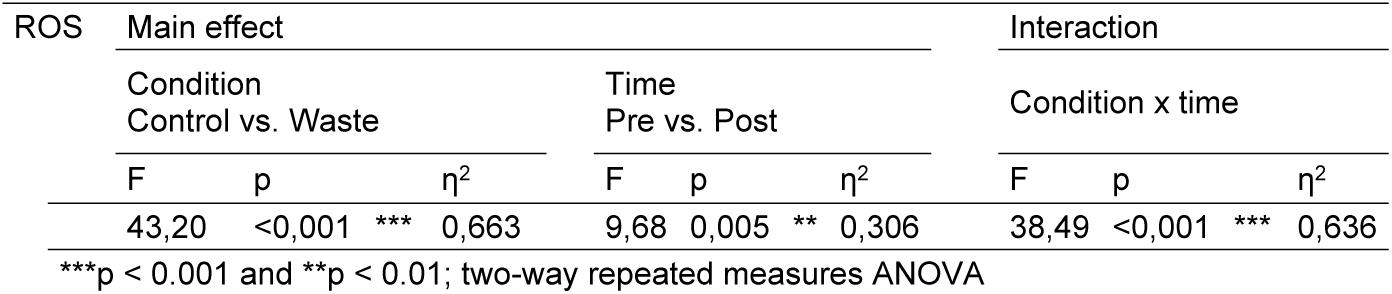
Results of two-way repeated measures ANOVA for the Restorative Outcome Scale (subjective restorativeness).

**Table 8.**
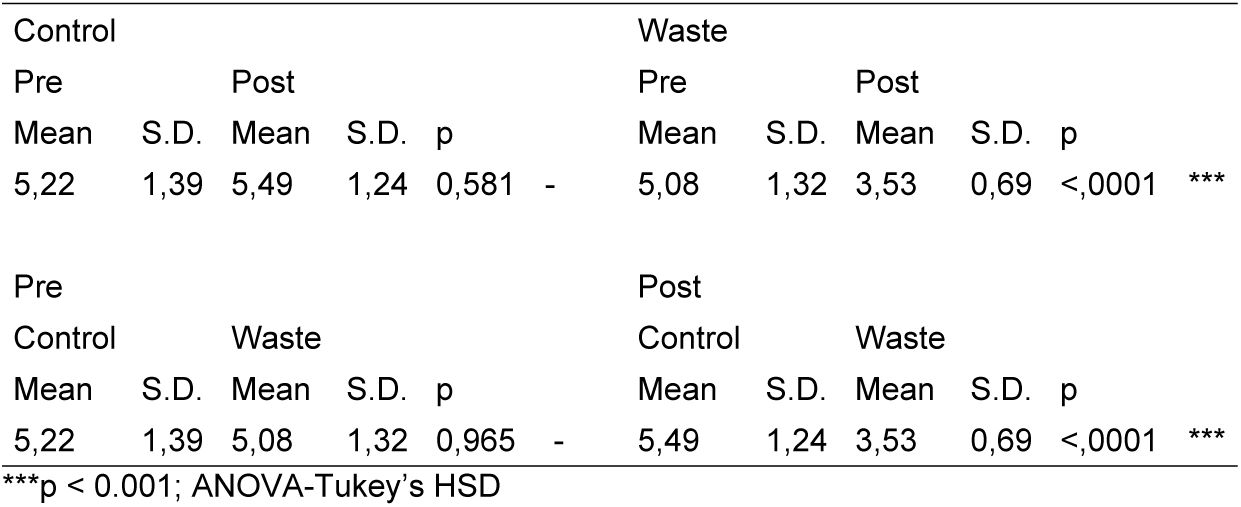
Results of the multiple comparison test between the control and experimental (setting) and pre-post (viewing forest without or with waste) for the Restorative Outcome Scale (subjective restorativeness).

In the case of the SVS, a two-way repeated measures ANOVA was used to investigate subjective vitality treating ‘Conditon’ and ‘Time’ as factors, and the interaction between these two factors was also assessed. The effects and interaction were significant (Table 9). The results of the multiple comparison Tukey’s HSD test (Table 10) showed that in the control forest, there was a significant increase in SVS scores after viewing the forest environment (Control: Pre vs. Post). There was no difference between the control forest and the forest with waste in the pretest (Pre: Control vs. Waste). A significant decrease of the SVS scores was observed after comparing the situation before viewing the forest with waste with that after viewing the forest with waste (Waste: Pre vs. Post); and when comparing the posttests in the control forest and in the forest with waste, the latter achieved significantly lower values (Post: Control vs. Waste).

**Table 9.**
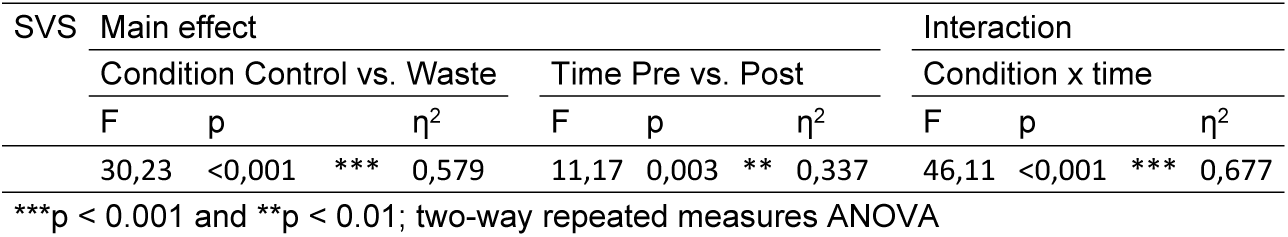
Results of the two-way repeated measures ANOVA for the Subjective Vitality Scale (subjective vitality).

**Table 10.** Results of the multiple comparison test between the control and experimental (setting) and pre-post (exposure to viewing forest) for the Subjective Vitality Scale (subjective restorativeness).

| Control |  |  |  |  | Waste |  |  |  |  |
| --- | --- | --- | --- | --- | --- | --- | --- | --- | --- |
| Pre |  | Post |  |  | Pre |  | Post |  |  |
| Mean | S.D. | Mean | S.D. | p | Mean | S.D. | Mean | S.D. | p |
| 4,91 | 1,03 | 5,40 | 1,13 | 0,031 * | 4,88 | 1,21 | 3,50 | 0,72 | <,0001 *** |
| Pre |  |  |  |  | Post |  |  |  |  |
| Control |  | Waste |  |  | Control |  | Waste |  |  |
| Mean | S.D. | Mean | S.D. | p | Mean | S.D. | Mean | S.D. | p |
| 4,91 | 1,03 | 4,88 | 1,21 | 0,999 - | 5,40 | 1,13 | 3,50 | 0,72 | <,0001 *** |

\*\*\*p < 0.001 and \* p < 0.05; ANOVA-Tukey's HSD

## Discussion

Forest bathing is an experience that unquestionably positively influences the psychological relaxation of people [1,4–6,8,10,11,13,13,14,21,23,40–44,44,45]. Consistent with previous studies [28,29], this study confirmed that some factors might interrupt this experience that is frequently called a restorative experience. Regarding all four scales used in the current research (POMS, PANAS, ROS, and SVS) and the ten total indicators of different psychological constructs, it might be concluded that each of them significantly reacts to the interruptive effect of an illegal open forest dump occurring in the forest environment. Negative indicators that are commonly described as signs of bad mental health, like Tension-Anxiety, Depression-Dejection, Anger-Hostility, Fatigue, Confusion, and the PANAS Negative, significantly increased after viewing the forest with waste compared to the pretest and control. In contrast, positive indicators describing good mental health, like Vigor, the PANAS Positive, the ROS, and the SVS, significantly decreased after exposure to the open dump. This is the evidence from the one case study in one forest stand and in one day; however, the study was conducted with a rigorous experimental procedure using a randomized, controlled crossover design. Based on this knowledge, it might be hypothesized that open dumps in a forest can lower the psychological relaxation of viewers and, therefore, this might influence their restorative experience during walks throughout the forest environment.

The rigorous usage of a crossover experiment and pretest without visual stimulation allowed us to deduce that viewing waste in the forest is enough to decrease the psychological relaxation of young adult visitors. The mechanism of these observations is hard to explain, and it is only known at this stage that some negative effects occurred and were observed in the results. There are two premises that might explain the occurrence of these effects. The first is the theory of identity with place or aspect, known as place identity [46]. According to that theory, participants might not definitively identify with open dumps, which lowers their restoration and increases the negative indicators. In a group of two hundred students, the authors prove [46] that a local environment induced more psychological relaxation than a non-local one. The second theory is associated with the biophilia hypothesis. According to this hypothesis, humans are biophilic; thus, they have an essential predisposition to stay in nature since it makes them feel good and appropriate [47]. An open dump in the forest is the opposite of these environments, and so a decrease in psychological relaxation is observed in this environment. However, the occurrence of waste in the forest is an interrupting factor that might influence the restorative experience of forest visitors. The explanation of this phenomenon needs further rigorous research.

These relevancies are new since the available literature includes no information about experiments testing the influence of illegal open waste in a forest environment on the psychological relaxation of participants. Therefore, until this research, there is nothing known about this effect, but there are some examples in the literature of other experiments testing some interventions in the forest on participants. One example is testing the effect of the slight thinning of a managed coniferous forest on landscape appreciation and psychological restoration [45]. In this experiment, the results were ambiguous. ‘Fatigue’ significantly increased after viewing the forest after thinning (p = 0.017) but ‘Confusion’ decreased significantly (p = 0.000); the other results were not significant when comparing the two conditions. In other research, the age of a stand was a disturbing factor, and greater restorativeness occurred from a pristine forest stand with old age than a young forest [48]. In another study, researchers evaluated the usage of a computer in a natural environment as a disturbing factor, and concluded that the usage of this equipment reduced psychological relaxation [28]. Also, viewing a forest with a disturbing factor such as a building decreased psychological relaxation [29].

Another example is the assessment of the impact of viewing vs. not viewing a real forest on psychological responses [49]. In this research, the authors investigated the impact of tents with enclosed/open sheets as a moderator when viewing a forest stand. Viewing a forest significantly decreases Tension-Anxiety, Depression, Fatigue and Confusion (p = from 0.031 to 0.001). In the current study, for participants viewing a forest after wearing opaque eye patches fifteen minutes before doing so, only one psychological indicator – the subjective vitality – was significantly increased. In another research conducted during the spring, other negative indicators decreased significantly. Also, most values of the negative psychological indicators did not decrease significantly in comparison to the control, but subjective vitality significantly increased after exposure to the forest environment [23]. The possible explanation of these results might be the disturbing factors of leaves and insects that occurred in the forest during the spring. Thus, significant changes in the case of only one from ten measured psychological indicators during forest bathing in the spring in North-East Poland are not disappointing because the results are almost consistent with previous studies.

It is also worth considering that factors that occur in a physical environment might influence the psychological relaxation of participants. Some environmental conditions were measured in this research, but only one – relative humidity – was significantly different in the forest with waste than in the control forest. The humidity is significantly, negatively correlated with mood disturbances (*r* = – 0.30) [13], but in the current research, the humidity did not have a significant effect. Furthermore, it was not a deciding factor in lowering the effects of negative psychological indicators because the values of the relative humidity were higher in the case of the forest with waste and increased negative values of the psychological response were still observable.

It was possible to stimulate participants’ olfactory or odor senses, which were not controlled in this experiment, but visual stimulation is probably the most important. Nevertheless, the participants were standing (during pretest) at some distance from the waste, and thus the olfactory effect was eliminated. During the experiment, none of the participants reported experiencing the olfactory effect of open waste. It is also not known what the real effect of an open dump in the forest is during normal everyday experiences; thus, it needs further investigations to test this effect during walking in the forest.

Besides, it is important to recognize that people experienced some kind of negative mood while walking and recreating in the forest with open dumps, even if it in an experimental situation and not a real forest. Also, the influence on people that clean these areas with waste is interesting. Nevertheless, these questions are left for further investigations.

## Conclusions

This study clarifies that the view of an open dump in a forest environment (in comparison to an undisturbed, control forest) might influence the psychological relaxation of respondents. What emerges is increased negative indicators of psychological relaxation and decreased positive indicators. In this study, it is also clarified that viewing the open dump in the forest environment has a negative effect on respondents, which is proved in a randomized, controlled crossover experiment. The explanation of this effect needs further investigations. Also, the effect of relaxation in a forest during spring was relatively small in these experiments. Only subjective vitality was stimulated by the control forest environment. In contrast, the olfactory stimulation and humidity in the experiment did not influence the expected negative effect of open dumps.

## Acknowledgements

Thanks to Robert Rydzewski for great technical assistance in organizing the experiment.

